# Histone chaperones exhibit conserved functionality in nucleosome remodeling

**DOI:** 10.1101/2022.01.13.476140

**Authors:** P. Buzón, A. Velázquez-Cruz, K. González-Arzola, A. Díaz-Quintana, I. Díaz-Moreno, W.H. Roos

## Abstract

Chromatin homeostasis mediates some of the most fundamental processes in the eukaryotic cell. In this regard, histone chaperones have emerged as major regulatory factors during DNA replication, repair, and transcription. However, the dynamic nature of these processes has severely impeded their characterization at the molecular level. Here we apply single-molecule probing by fluorescence optical tweezers to follow histone chaperone dynamics in real-time. The molecular action of SET/template-activating factor-Iβ and nucleophosmin 1—representing the two most common histone chaperone folds—were examined using both nucleosomes and isolated core histones. We show that these chaperones present binding specificity for partially dismantled nucleosomes and are able to recognize and disrupt non-native histone-DNA interactions. Furthermore, we reveal that cytochrome *c* inhibition of histone chaperones is coupled to chaperone accumulation on DNA-bound histones. Our single-molecule approach shows that despite the drastically different structures of these chaperones, they present conserved modes of action mediating nucleosome remodeling.

## Introduction

The nucleosome is the minimal unit of structural organization in eukaryotic genomes^1^. It comprises ~146 base pairs (bp) of DNA that wraps around an octameric protein complex involving two copies of each core histone (H2A, H2B, H3, and H4)^2^. This supramolecular arrangement provides genome stability and confinement, while serving as the ultimate regulatory barrier that mediates DNA accessibility. Thus, nucleosome dynamics plays a unique role in many of the most fundamental cellular processes—from genome replication and repair to gene expression. Nucleosome homeostasis is highly regulated, and therefore coordinated by the action of many different nuclear factors, including histone-modifying enzymes^3^, ATP-dependent chromatin remodelers^4^, and histone chaperones^5^. In addition to the nuclear machinery, other cellular factors have been identified to play critical roles in diverse processes associated with chromatin remodeling. Particularly, the mitochondrial protein cytochrome *c* (C*c*) has been recently reported to be translocated into the nucleus in the context of DNA damage, where it acts as a histone chaperone inhibitor^6, 7^.

In this framework, histone chaperones have emerged as the main factors of the cellular machinery responsible for regulating histone availability^8^. These chaperones are often multifunctional proteins involved in many processes related to histone metabolism, such as folding, oligomerization, transport, deposition and eviction, storage, post-translational modifications, and nucleosome assembly^5, 9^. In general, histone chaperones perform their tasks by direct association with single and oligomeric histones, mediating specific interactions and preventing aggregation. Interestingly, histone chaperones also modulate antagonistic processes: histone-DNA deposition and eviction. However, our understanding of the mechanisms behind these chaperoning processes is still limited. Moreover, histone chaperone activities are found in a wide range of different protein families, which show little sequence similarities between them. Nevertheless, two characteristic structural folds have been identified for their specific role as histone chaperones: the dimeric Nucleosome Assembly Protein 1-like (NAP1-like) fold and the pentameric nucleoplasmin fold^9^.

The study of histone chaperone activities has proven experimentally challenging due to the intrinsic instability of histones and their aggregation propensities, particularly in the presence of DNA. Here, we present a set of single-molecule manipulation strategies, combining optical tweezers (OT) and confocal fluorescence microscopy (CFM)^10, 11^, to investigate chaperone activity in real time with molecular resolution. We selected two chaperones that represent the two most conserved histone-chaperone folds: SET/template-activating factor-Iβ (SET/TAF-Iβ) that belongs to the NAP1-like family^12^, and nucleophosmin 1 (NPM), which exhibits the nucleoplasmin fold^13^ (Fig. S1). Specifically, we probed the chaperone activity of both SET/TAF-Iβ and NPM acting on individual histones, as well as in the context of the nucleosome. Chaperones showed binding specificity for partially disrupted nucleosomes, suggesting histone exposure as a key factor during nucleosome recognition. Moreover, the relative decrease in histone-DNA affinity in the presence of chaperones was characterized through kinetic measurements, while providing direct observations on histone shielding and eviction. Finally, histone eviction assays in the presence of C*c* revealed that the inhibition of the chaperoning activity is coupled to the accumulation of chaperones on DNA-bound histones, suggesting new mechanisms for chaperone activity regulation. Hereby our results provide new molecular insights into the regulatory functions of histone chaperones coordinating histone-DNA interactions.

## Results

### Both SET/TAF-Iβ and NPM exhibit specificity for partially unwrapped nucleosomes

Histone chaperones are known to mediate various processes involved in the regulation of histone-DNA interactions. One of their main functions is to assist nucleosome assembly and disassembly through histone deposition and eviction. To investigate these highly dynamic processes with molecular resolution, we employed a combination of OT and CFM. OT allows monitoring the mechanical unwrapping of reconstituted nucleosomes by performing force-extension curves (FECs) on individual DNA molecules, while CFM provides direct visualization of fluorescently labeled chaperones and histones with single-molecule resolution.

Nucleosomes were *in vitro* reconstituted using ~8.3 kbp DNA molecules lacking artificial nucleosome positioning sequences, and isolated by using OT and microfluidics. Subsequently, individual molecules of DNA containing nucleosomes were brought to a solution of either SET/TAF-Iβ or NPM, where nucleosomes were incubated for a minute before being mechanically unwrapped by pulling on the DNA (Fig. 1a). A representative FEC is shown in Fig. 1b, displaying a typical saw-tooth pattern during tether extension (grey) resulting from nucleosome unwrapping. Retraction curves (green) also showed abrupt changes in force, although to a lesser extent, indicative of DNA rewrapping. We use the extensible worm-like chain (eWLC) model to fit the smooth regions of the FEC (dashed lines in Fig. 1b) and determine the different values of apparent contour length (*L_c_*) of the tether (see methods). Then, from the values of *L_c_*, the changes in apparent contour length (Δ*L_c_*) after every rupture event of the curve are calculated, corresponding to the number of unwrapped base pairs (Fig. 1b, c). Our nucleosome unwrapping experiments reported a major Δ*L_c_* population at 72 ± 5 bp (center ± SD) (Fig. 1c), in agreement with the values reported in the literature for the two-step nucleosome unwrapping mechanism^14-21^. In addition, a smaller and wider peak was found at ~30 bp (Fig. 1c). This small population could be due to partial nucleosome unwrapping/rewrapping multistep events, smaller than the canonical ~72 bp. Also, changes in nucleosome reorientation with respect to the direction of the applied force could be taking place, as previously proposed^18^.

**Fig. 1.**
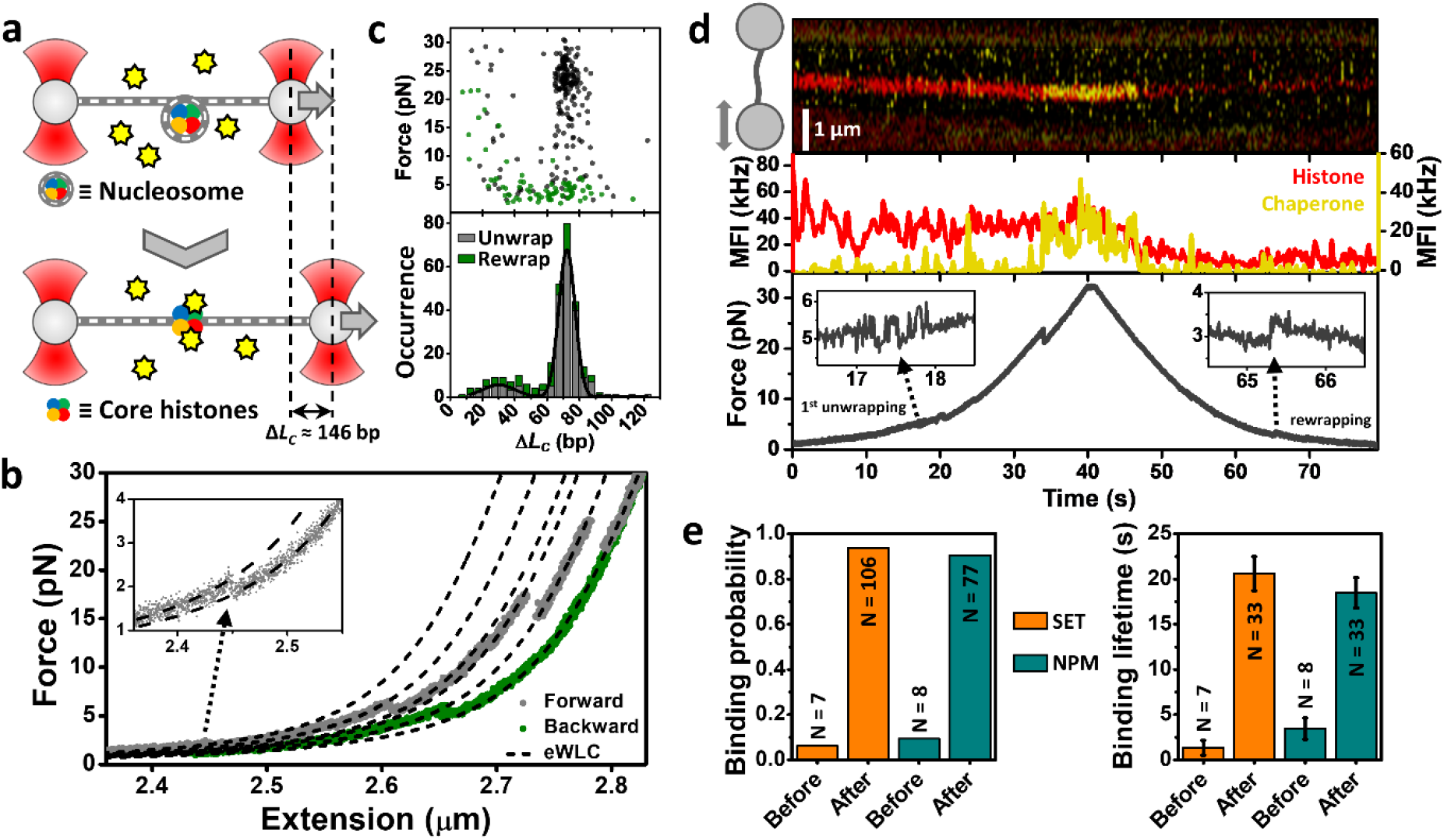
Chaperone-histone interactions in the context of the nucleosome. **a,** Scheme of a nucleosome unwrapping experiment. Nucleosomes were mechanically unwrapped in the presence of 1-2 nM concentration of chaperone. Δ*L_c_* shows the expected total change in contour length upon unwrapping a single nucleosome. **b**, Representative FEC showing several unwrapping events during the forward curve (dark grey) and rewrapping during the backward curve (green). Inset, zoom in on the first unwrapping event of the forward curve. Black dashed lines represent eWLC fits to the forward curve (see Methods). **c**, Lower panel, stacked bar histogram of the Δ*L_c_* values obtained from the forward (N = 209; dark grey) and backward (N = 81; green) FECs. The fitting of the unwrapping event distribution to a double Gaussian reported 72 ± 5 bp and 30 ± 10 bp (center ± SD). Upper panel, force distribution at which the changes in *L_c_* occurred. **d**, Correlated FEC and fluorescence imaging showing histone eviction upon nucleosome unwrapping. Upper panel, kymograph recorded at 4 Hz of a fluorescently labeled nucleosome (red) and SET/TAF-Iβ (yellow) during the FEC. The region occupied by the beads has been darkened for visualization purposes. Middle panel, mean fluorescence intensity (MFI) obtained from the traces shown in the upper panel, histones (red; left axis) and chaperone (yellow; right axis). Lower panel, force *versus* time plot of the FEC. Insets highlight the un/rewrapping events found at low forces. **e**, Left panel, probabilities obtained for SET/TAF-Iβ (orange; N = 113 binding events) and NPM (dark green; N = 85) binding to histones before and after an unwrapping event was detected. Right panel, dwell time of the binding events identified before and after any unwrapping event, represented as mean ± SEM. N = 40 (SET/TAF-Iβ) and N = 41 (NPM) binding events. The mean binding lifetimes obtained after unwrapping should be considered as lower limits of the real values, as some chaperones remained bound until the end of the experiment.

In addition to the *L_c_* changes, fluorescently labeled histones (red, Fig. 1d) and chaperones (yellow) were directly visualized by two-color CFM. Nucleosome labeling did not report any differences in Δ*L_c_* when compared to FECs of non-labeled nucleosomes (Fig. S2a). We spotted two major actions when looking at the histone fluorescence signal in combination with force-extension experiments. Histones were found to either remain bound to DNA, or unbind after nucleosome unwrapping (Fig. S2b, c); consistent with the lower number of rewrapping events with respect to unwrapping (Fig. 1c). Moreover, fluorescent experiments also revealed that in the absence of histones, using bare DNA molecules, chaperones do not interact with DNA, as discussed in the next section.

By correlating the FECs with the fluorescence imaging, we were able to reveal, in real time, the action of chaperones during histone eviction from partially unwrapped nucleosomes. Fig. 1d shows how the SET/TAF-Iβ signal (yellow) colocalizes with the histone signal (red) for ~15 s followed by a correlated decrease in both signal intensities. Not all fluorescently labeled histones within the nucleosome are evicted by the chaperone, as perceived by the remaining red fluorescence intensity (Fig. 1d). Eviction is identified by the drop of the chaperone signal (yellow) to background levels. According to our brightness calibration (Fig. S3), a fluorescently labeled SET/TAF-Iβ protein presents ~20 kHz (photons•10^3^/s) in mean fluorescence intensity (MFI). The dimeric nature of SET/TAF-Iβ implies that on average every dimer displays two fluorophores (see Methods), allowing us to identify a drop of ~20 kHz as chaperone unbinding, as two fluorophores typically do not bleach simultaneously. Hence, Fig. 1d shows an example capturing histone eviction carried out by an individual chaperone. Another example of eviction where histone signal dropped to background levels is shown in Fig. S2d. Moreover, we also find examples where chaperones could either remain bound until the end of the FEC or unbind without any change in the histone signal (Fig. S2e). The latter observation might be strongly conditioned by our nucleosome labeling procedure (see Methods). Interestingly, these experiments also revealed that both chaperones have a clear preference binding to partially disrupted nucleosomes (Fig. 1e). More than 90% of the chaperone binding events were found after unwrapping events, suggesting that histone exposure greatly favors the recognition of nucleosomes by these histone chaperones. This observation was further supported by the analysis of the binding lifetimes of both chaperones (Fig. 1e). The binding events identified after partial or complete nucleosome unwrapping were significantly longer, indicative of the higher affinity of chaperones for exposed histones.

### Histone chaperones are recruited to DNA by DNA-bound histones

To gain a better understanding of the mechanisms underlying the various observations described above, including chaperone-histone interaction and eviction, we continued studying the chaperone activity of SET/TAF-Iβ and NPM with single histones. DNA molecules (~48 kbp; see methods) were isolated using OT and microfluidics and incubated in a solution containing either of the individual core histones (H2A, H2B, H3, or H4) to allow the formation of histone-DNA complexes. Subsequently, these complexes were brought to a solution of fluorescently labeled histone chaperones to monitor their interaction (Fig. 2a). While control experiments without histones do not report any chaperone binding event, measurements including histones confirm that chaperones are recruited to DNA by DNA-bound histones. Fig. 2b shows the presence of chaperone-histone-DNA complexes and their evolution during the first 5 min of incubation in the chaperone solution. CFM imaging reveals how the fluorescence signal at certain regions of the DNA molecule disappear during the incubation time, indicative of chaperone unbinding (Fig. 2b, c). In addition, the overall signal measured as MFI reported that the major changes in fluorescence occur during the first 2-3 min (Fig. 2b, right panel). All core histones behaved similarly and showed the same trend; for both SET/TAF-Iβ and NPM the fluorescence signal disappeared partially, but not completely, within 5 min (Fig. S4a).

**Fig. 2.**
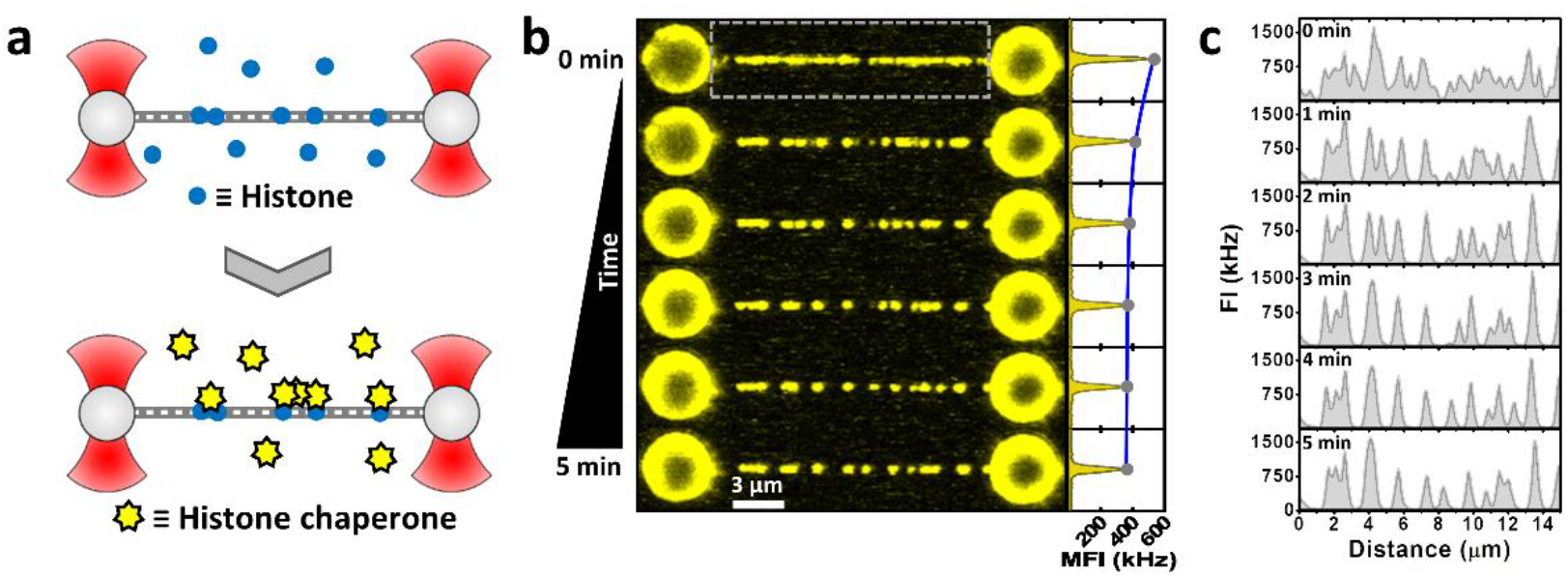
Direct visualization of histone recognition by chaperones. **a,** DNA molecules are stretched to a fixed distance, corresponding to 10 pN tension, and then incubated in a solution of individual core histones at 100 nM concentration for ~30 s. Next, histone-DNA complexes were brought to a solution of fluorescently labeled histone chaperones at 5 nM concentration, and observed for 5 min. **b**, Left panel, fluorescence images recorded at one-image-per-minute frequency (image time ≈ 5 s) during the incubation of a H2B-DNA complex in the presence of 5 nM NPM. Right panel, MFI measured from the area between the beads (dashed rectangle) at each time point. **c**, Fluorescence intensity (FI) values corresponding to the signal of a single line scan of one-pixel thickness along the DNA molecule shown in (b) at different time points.

By continuously scanning along the DNA molecule at much higher frequency we were able to capture the truly dynamic nature of the process (Fig. S4b). We observed a fast binding of chaperones, followed by a clear decrease in fluorescence intensity within 3 min. As chaperones do not bind to bare DNA under our conditions, these results suggest that the number of DNA-bound histones decreases over time upon incubation with histone chaperones, supporting our observations of histone eviction in the context of mechanically unwrapped nucleosomes.

### Histone chaperones prevent DNA bridging through histone shielding and eviction

To further study histone-DNA unbinding kinetics, we applied a strategy based on measurements of DNA de/condensation upon protein un/binding. This assay, which is independent of fluorescence intensity measurements, allows monitoring histone binding with higher temporal resolution. DNA molecules held in the presence of histones experienced an increase in tether tension upon binding, and hence, a decrease in the bead-to-bead distance (Fig. S5a). Both tether tension and bead-to-bead distance gets partially restored to its original, that of bare DNA, when histone-DNA complexes are brought into the chaperone solution (Fig. 3a and S5a). Fig. 3a shows averaged kinetic traces for all individual histone-DNA complexes upon incubation with either NPM (dark green), SET/TAF-Iβ (orange), or buffer (grey) as control.

**Fig. 3.**
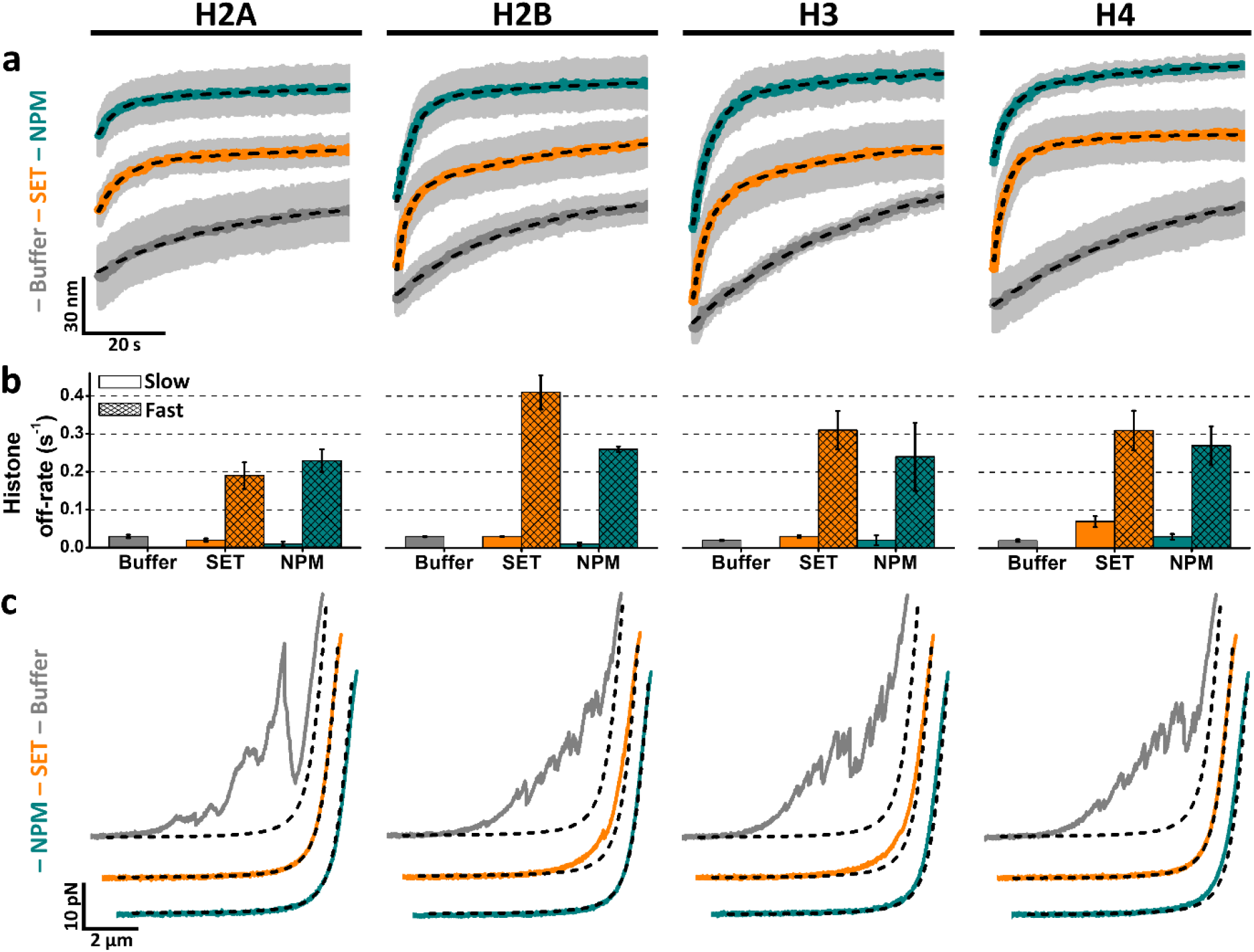
Eviction and shielding of DNA-bound histones mediated by histone chaperones. **a,** Averaged kinetic traces of bead-to-bead distance over time for histone-DNA complexes during their incubation against buffer (grey), 5 nM SET/TAF-Iβ (orange), and 5 nM NPM (dark green). Averaged curves were generated from individual traces (N ≥ 5) and fitted to either a single or double exponential function (black dashed line) for the first 2 min of incubation. For visualization purposes only the first minute is shown. Grey shades represent SEM. **b**, Off-rates obtained from the fits presented in (a). Traces obtained in buffer could be fitted to a single exponential, while chaperone traces presented two off-rates. Error bars represent SEM. **c**, Representative FECs obtained after 5 min incubation in buffer (grey), 5 nM SET/TAF-Iβ (orange), and 5 nM NPM (dark green). Black dashed lines represent simulated bare DNA curves generated with the eWLC model and experimentally measured parameters (see methods).

Histone-DNA complexes that were brought to the buffer solution showed a monotonic increase in bead-to-bead distance reporting similar off-rates, between 0.02 and 0.03 s^-1^ (Fig. 3a, b). However, in the presence of chaperone, significantly faster kinetics were found in all cases. These traces showed a bimodal growth composed of a slow component, which resembles the off-rates measured in buffer, and a second kinetic component approximately 10 times faster (Fig. 3a, b). These results are in agreement with the behavior revealed by CFM experiments, in which the amount of bound chaperones decreased over time (Fig. 2b, c). In addition, this kinetic analysis indicates that histones are being actively removed, evicted, from DNA by the action of histone chaperones. To further validate this, we used a truncated version of SET/TAF-Iβ (SET/TAF-Iβ-ΔC), lacking the C-terminal disordered acidic domain and displaying a lower affinity for histones^12^. As could be anticipated, kinetic traces in the presence of SET/TAF-Iβ-ΔC showed a clear decrease in the histone eviction rate when compared with full-length SET/TAF-Iβ (Fig. S5b, c).

We then attempted to gain further insights into the chaperone-histone complexes that remained bound to DNA (Fig. 2b, c and S4). Thus, FECs of histone-bound DNA were performed after 5 min incubation in either buffer or chaperone solution. FECs without chaperone displayed a large amount of DNA loops, generated during the relaxation of the tether tension, and disrupted during pulling (Fig. 3c). These DNA loops are most likely formed through protein-DNA interactions between distant regions of the DNA molecule, and nonspecific protein-protein interactions. Interestingly, when FECs were performed in the presence of either of the chaperones, the curves turned substantially smoother, in some cases resembling bare DNA curves (Fig. 3c). These experiments, combined with CFM imaging (Fig. 2b and S4), prove the capabilities of both SET/TAF-Iβ and NPM to accumulate on DNA-bound histones and shield nonspecific histone-histone and histone-DNA interactions, preventing DNA bridging.

Ultimately, our results unveil, at the molecular level, two major regulatory functions of chaperones in the context of chromatin remodeling. Both SET/TAF-Iβ and NPM showed the ability to efficiently disrupt noncanonical histone-DNA interactions through histone removal. When histone eviction cannot be achieved, histone shielding is used to prevent the formation of further nonspecific interactions, thus ensuring genome integrity.

### Cytochrome *c* inhibits histone chaperone activities

Cytochrome *c* (C*c*) is a multifunctional^22^ mitochondrial protein known to play a major role in the electron transport chain^23^, a key process in the synthesis of ATP. In addition, C*c* has been shown to be released from mitochondria and to translocate into the nucleus under DNA damage conditions^6, 24-27^. Although the nuclear role of C*c* is not fully understood, it has recently been shown that following DNA breaks, C*c* specifically interacts with SET/TAF-Iβ^6^, and NAP1-Related Protein 1 (NRP1)^7^, a plant homologous of SET/TAF-Iβ, acting as a histone chaperone inhibitor. Therefore, we decided to include C*c* in our single-molecule kinetic experiments to address the physiological relevance of the chaperone activities reported in Fig. 3.

Due to the very similar behavior exhibited by all core histones, kinetic experiments were performed only with histone H3, measuring SET/TAF-Iβ and NPM chaperone activity at increasing concentration of C*c* (Fig. 4a, b). Explicitly, H3-DNA complexes were brought to a mixture solution of chaperone and C*c*, keeping chaperone concentration constant (5 nM), while varying C*c* concentration. We found a clear decrease in the fast kinetic component with increasing C*c* concentrations, proving that histone eviction activity is weakened by C*c* (Fig. 4a, b). Particularly, chaperone activity was completely inhibited at 0.5 μM (SET/TAF-Iβ) and 2 μM (NPM) concentration of C*c*. The same behavior was observed with SET/TAF-Iβ-ΔC (Fig. S5d). Control experiments performed with C*c* at these saturating concentrations, but without chaperones, did not alter the H3 intrinsic unbinding rate (Fig. S6b). The inhibition of histone shielding could not be addressed as C*c* interacts with DNA and promotes the formation of DNA loops. Similarly, nucleosome unwrapping could not be detected in the presence of C*c* as the disruption of these DNA loops showed much greater signals than the ones obtained from nucleosome disruption. Performing FECs in the absence of histones revealed that C*c* both alone and in a mixture with any of the abovementioned chaperones, promotes DNA bridging (Fig. S6).

**Fig. 4.**
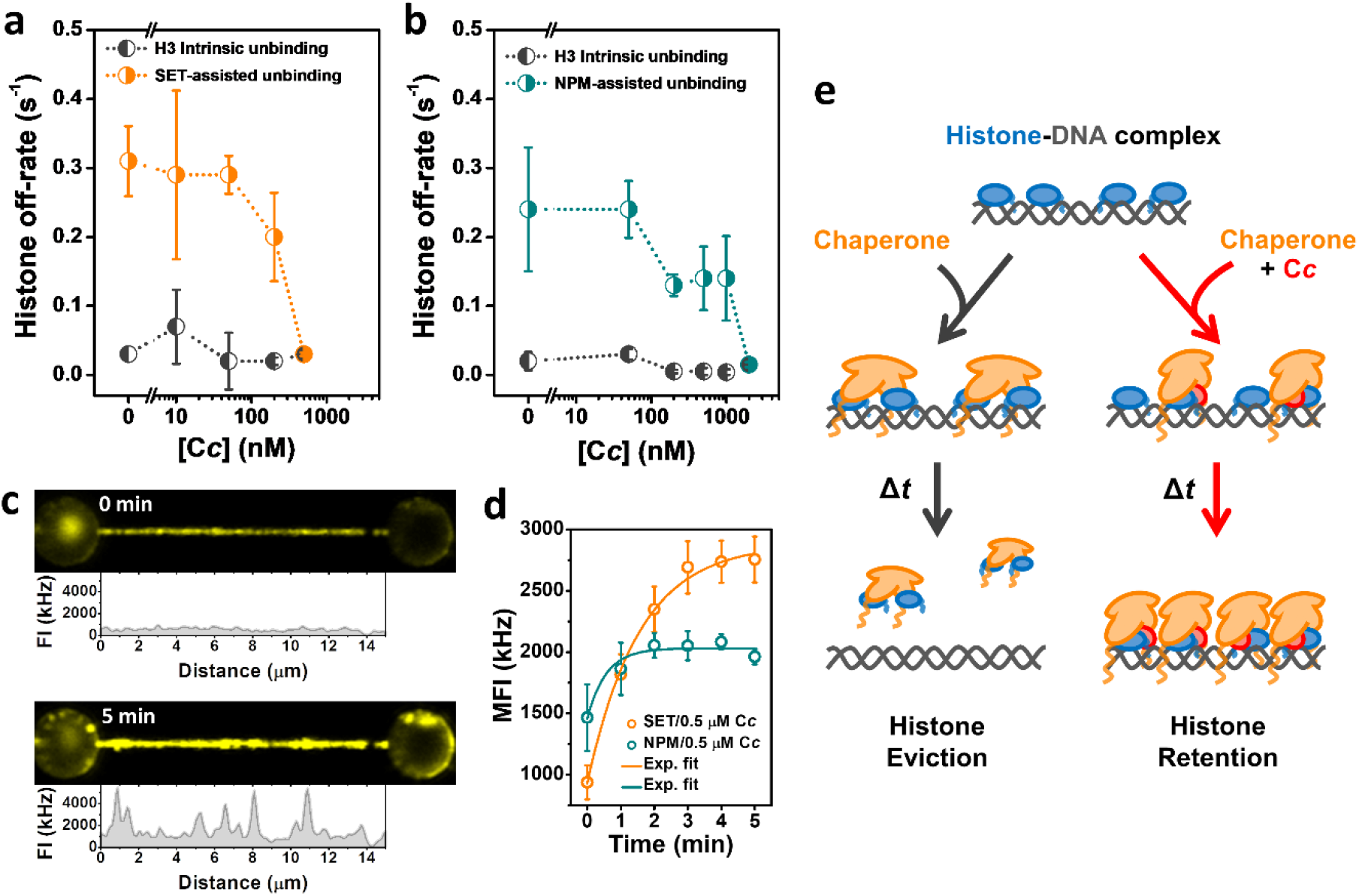
Inhibition of histone chaperone activity by C*c*. **a**, Off-rates of H3 dissociation from DNA as a function of C*c* concentration (10-500 nM) and in the presence of 5 nM SET/TAF-Iβ, as obtained from averaged kinetic traces. The slow (H3 intrinsic unbinding) rate is depicted in black, while the fast (SET/TAF-Iβ assisted unbinding) rate is colored in orange. Error bars represent SEM. **b**, Off-rates obtained as described in (a) for a solution comprising 5 nM NPM and varying concentrations of C*c* (50 nM-2 μM). NPM-assisted unbinding is colored in dark green and the intrinsic H3 unbinding rate in black. Error bars represent SEM. **c**, Representative fluorescence images of a H3-DNA complex in a solution of 5 nM SET/TAF-Iβ and 500 nM C*c* at 0 and 5 min incubation. FI values correspond to the signal of an individual line scan along the DNA molecule. **d**, MFI over time obtained from CFM images as the fluorescence images shown in (c), for SET/TAF-Iβ (orange; N = 5) and NPM (dark green; N = 3), both with 500 nM of C*c*. Error bars represent SEM. Fits to an exponential function (solid lines) reported 0.64 ± 0.07 s^-1^ (SET/TAF-Iβ) and 2 ± 1 s^-1^ (NPM). **e**, Schematic representation of C*c*-mediated inhibition of histone eviction by chaperones. Histone chaperones (orange) facilitates the eviction of individual core histones (blue) from DNA. However, chaperone-assisted histone eviction becomes hindered and eventually halted when an excess of C*c* (red) is present. Furthermore, chaperones accumulate onto DNA, presumably interacting with both histone and C*c*.

In addition to the kinetic analysis, CFM imaging of fluorescently labeled chaperones in presence of C*c* also pointed toward a reduction in histone removal activity. Fig. 4c shows how C*c* promotes chaperone accumulation onto DNA upon incubation, reverting the behavior described previously for chaperones alone, where fluorescence intensity would decrease over time (Fig. 2b, c). Chaperone-C*c* localization on DNA showed a monotonic increase over the first minutes of incubation before reaching saturation (Fig. 4d). We report chaperone accumulation rates of 0.64 ± 0.07 s^-1^ (SET/TAF-Iβ) and 2 ± 1 s^-1^ (NPM), both at 500 nM of C*c*. Consequently, chaperones appear to be still able to interact with histones even when C*c* is present, while histone eviction activity is inhibited. Although control experiments carried out without histones showed that C*c* has also the ability to bind to DNA and recruit chaperones (Fig. S6), we found that chaperones colocalize on DNA to a much lesser extent than in the presence of histones (Fig. 2b, c). Finally, these results reveal what seems to be a very specific function of C*c* during chaperone activity regulation, which involves the inhibition of histone eviction while promoting chaperone accumulation; a function that could potentially be conserved for many other histone chaperones (Fig. 4e).

## Discussion

Correlated OT and single-molecule fluorescence measurements revealed that both SET/TAF-Iβ and NPM are recruited to DNA by histones. Previous quantification of the binding affinity of SET/TAF-Iβ and NPM for histones in absence of DNA provided dissociation constants (*K*_D_) in the micromolar concentration range by Isothermal Titration Calorimetry^6^. Although these histone chaperones have also been described to exhibit significant DNA binding capabilities^12, 28, 29^, we have shown that in the low nanomolar range they specifically colocalize onto DNA only if histones are present. Transient binding of chaperones to DNA might still occur, but our experiments prove that those must be in the order of ~200 ms or shorter, according to the temporal resolution achieved by fluorescence microscopy. Our kinetic analysis of histone-DNA interactions reported off-rates between 0.02 and 0.03 s^-1^. These values represent the kinetic constant (*k_off_* for individual histones under moderate ionic strength conditions (50 mM KCl). Assuming a *k_on_* between 10^8^-10^6^ M^-1^s^-1^, our results suggest high affinity between histones and DNA, with *K*_D_ values between the picomolar and low nanomolar range. Affinities that are consistent with estimations of the DNA binding properties of histone octamers^30-32^. Importantly, these affinities decrease approximately ten times when 5 nM of chaperone is added to the solution (Fig. 3a, b), proving SET/TAF-Iβ and NPM histone eviction mechanisms. Histone eviction activity was further visualized by CFM, monitoring the decrease of bound chaperones over time. However, core histones are known to arrange as heterodimers, H2A-H2B and H3-H4, or tetramers in the case of (H3-H4)2, prior to their deposition on DNA^33^. Therefore, our observations quantitatively describe the decisive ability of these chaperones to recognize non-canonical histone-DNA interactions, where histones are not in their dimeric arrangement, and trigger histone eviction. These findings are in line with the reported NAP1 activity mediating the interaction between H2A-H2B dimers and DNA^34^.

Fluorescence imaging of chaperone-histone-DNA complexes also showed that not all histones are removed during chaperone incubation. These complexes were always found to be distributed as discrete high intensity spots (Fig. 2b, S4), instead of a homogeneous low intensity coverage of DNA molecules. Furthermore, our estimation of the count rate per fluorophore (Fig. S3) suggests that these complexes include tens, and sometimes hundreds, of chaperone molecules per diffraction limited spot. Hence, we hypothesize that the remaining chaperone-histone complexes are composed of histone aggregates that cannot be removed by the action of the chaperones. Nonetheless, both SET/TAF-Iβ and NPM displayed an additional mechanism to tackle these aggregates and prevent non-canonical histone-DNA interactions. Experiments probing DNA bridging (Fig. 3c) in combination with CFM imaging (Fig. 2b and S4) showed that these chaperones can gather around histone aggregates and efficiently shield their DNA interacting regions. Thus, chaperones avoid the formation of nonspecific DNA-histone interactions, and hence DNA bridging, which explains their capabilities to prevent genome condensation and preserve its integrity.

C*c* has been shown to act as a histone chaperone inhibitor of NAP1-like chaperones following DNA damage^6, 7^. Specifically, C*c* showed the capability to inhibit nucleosome assembly. Furthermore, C*c* was identified to interact with the histone binding site of those chaperones, suggesting its capacity to displace chaperone-histone complexes in the absence of DNA. Indeed, we have demonstrated that C*c* is able to completely suppress histone eviction (Fig. 4a, b). However, the inhibition of chaperone activity does not seem to fully block chaperone-histone interactions. On the contrary, fluorescence imaging showed the accumulation of chaperones on histone-DNA complexes (Fig. 4c, d). In addition, micromolar concentrations of C*c* alone display the ability to recruit chaperones to DNA (Fig. S6), although to a lesser extent than histones (Fig. 2b, c). Taken together, these observations seem to indicate a more complex inhibition mechanism that differs from the action of competition by identical binding sites (Fig. 4e). Our single-molecule experiments point towards the importance of DNA in the inhibitory activity of C*c*, where C*c*-DNA and the intrinsic chaperone-DNA affinities might be playing a major role. In addition, long-range allosteric effects have already been described for C*c* modulating the activity of another histone chaperone, the Acidic leucine-rich Nuclear Phosphoprotein 32B (ANP32B)^35^. Hence, we conclude that allosteric effects might also be taking place upon C*c*-chaperone interaction, which still allows chaperone-histone interactions while blocking chaperoning activity.

Histone eviction activity was also identified for nucleosome-forming core histones (Fig. 1d). However, our nucleosome unwrapping experiments described a far more complex behavior than the one observed with individual histones. Indeed, chaperones are known to mediate histone eviction as well as deposition of histone heterodimers. Chaperones showed the capability to either remain bound for periods of time that would exceed the length of the experiment, or unbind without any change in the histone signal. In addition, our histone labeling strategy prevented us from performing a quantitative analysis of this chaperone activity, nor to address questions related to chaperone specificity for H2A-H2B or H3-H4 dimers. Despite this, correlated fluorescence imaging and mechanical unwrapping experiments revealed the clear preference of these chaperones to interact with dismantled, or partially unwrapped, nucleosomes (Fig. 1e). These findings imply that neither SET/TAF-Iβ nor NPM would have the ability to recognize or destabilize nucleosomes in the absence of other remodeling factors. Interestingly, other chaperones have proven their capacity to destabilize nucleosomes without other factors: NAP1^36, 37^, FAcilitates Chromatin Transcription (FACT)^38^ and nucleolin^39^. Regarding the chaperones studied here, SET/TAF-Iβ is known to mediate chromatin decondensation^40^ through its interaction with the linker histone H1^41^, the chromatin remodeling protein prothymosin α^42^, and the transcription coactivator cAMP-Response Element Binding (CREB)-binding protein^43^. Moreover, NPM has also been identified to directly interact with histone H1^44^, promote acetylation-dependent chromatin transcription^45^, and mediate both nucleosome formation and chromatin decondensation^46^, among other cellular functions^47^. Therefore, our data suggest that the execution of these functions associated with SET/TAF-Iβ and NPM could be happening through cooperation with other cellular factors that would assist nucleosome recognition and/or nucleosome disruption.

In summary, this study has revealed a set of chaperone activities that seem to be conserved in the two most representative chaperone folds found in eukaryotes, the NAP1-like and nucleoplasmin-like folds. Both SET/TAF-Iβ and NPM exhibited specificity for partially unwrapped nucleosomes, as well as for DNA-bound histones in the absence of nucleosomes. Chaperones also showed the ability to efficiently frustrate non-nucleosomal interactions between histones and DNA, promoting histone eviction. Moreover, the uncovered features of the inhibitory mechanism exerted by C*c* indicated that chaperone activities can be suppressed without chaperone-histone interactions being fully hindered. Thus, our results evince that these histone chaperones share fundamental activities, mediating histone recognition in a very similar manner—potentially present in many other histone chaperones. Hereby, we have provided new molecular details into the mechanisms behind these processes mediated by histone chaperones as well as their regulation.

## Methods

### Protein Samples

The pET3a expression plasmids coding for core histones from *Xenopus laevis* were obtained from Dr Tim Richmond (Institute of Molecular Biology and Biophysics, Switzerland). Human NPM and SET/TAF-Iβ constructs were cloned in frame with an N-terminal 6xHis-tag in pET28a vectors. The DNA coding for human C*c* was in a pBTR1 plasmid^48^, along with the yeast heme lyase for proper protein folding.

For the fluorescent labeling of SET/TAF-Iβ and C*c* by maleimide derivatization, a cysteine was introduced in their sequences by site-directed mutagenesis. The point mutation Q69C was introduced in the full-length (SET/TAF-Iβ-Q69C) and the C-terminal deletion (SET/TAF-Iβ-ΔC-Q69C, amino acids 1-225) constructs. The C*c*-E104C mutant was produced as previously described^49^.

All proteins were expressed in *E. coli* BL21(DE3) electrocompetent cells and grown in Luria-Bertani medium. Core histones were purified following a simplified version of the protocol reported by Luger *et al*.^50^, with a single-step gravity-flow chromatography using a carboxymethyl cellulose resin (Whatman). The different histone chaperone constructs were purified by gravity-flow chromatography using a Ni-NTA resin (GE Healthcare), according to the manufacturer’s instructions. C*c*-E104C purification was performed by cation exchange chromatography, with a Nuvia S column (Bio-Rad) and using a fast protein liquid chromatography (FPLC) system (NGC Chromatography System Quest 10, BioRad).

The concentration of individual histones was estimated using the Bradford assay^51^, whereas C*c* was quantified in its reduced state by measuring its absorbance at 550 nm (ε_550_ = 28.92 mM^-1^ cm^-1^). All proteins were stored at −80 °C until use.

### Fluorescent labeling of histones chaperones

Both chaperones SET/TAF-Iβ-Q69C and NPM were fluorescently labeled using Alexa Fluor 532 C_5_ maleimide (ThermoFisher). NPM was labeled by means of its endogenous exposed Cys104. Cys reduced state was ensured by incubating the chaperones at low micromolar concentrations with 10 mM dithiotreitol (DTT) for 30 min on ice. DTT was removed by applying the mixture to a PD Minitrap G-25 column (GE Healthcare). Labelling reactions were carried out by adding 1:10 excess of fluorophore reagent to Cys residue and incubating for 2 hours at 4 °C. After incubation, the reaction was ended by adding 10 mM DTT, and the excess of dye was removed by using a PD Minitrap G-25 column for NPM or a Superdex 200 10/300 GL (GE Healthcare) size exclusion chromatography column for SET/TAF-Iβ. Final protein concentration and labeling efficiency were determined spectrophotometrically according to the guidelines provided by the manufacturer (ThermoFisher). Labeling efficiencies were ~95% and ~75% for SET/TAF-Iβ and NPM respectively, implying two fluorophores per dimeric SET/TAF-Iβ molecule and four fluorophores per NPM pentamer on average. Chaperone concentrations were determined spectrophotometrically at 280 nm (ε_280_ 16,960 M^-1^ cm^-1^ for NPM and ε_280_= 32,430 M^-1^ cm^-1^ for SET/TAF-Iβ and SET/TAF-Iβ-ΔC). NPM and SET/TAF-Iβ(-ΔC) concentrations were expressed in their pentameric and dimeric forms, respectively. Labeled proteins were flash frozen and stored at −80 °C.

### Nucleosome assembly and fluorescent labeling

Nucleosome reconstitution was performed by salt dialysis on biotinylated DNA (pKYB1 vector; New England Biolabs) without any nucleosome positioning sequence, similarly to previously reported protocols^52^. A mixture of *Xenopus Laevis* core histones at 75 nM concentration (each histone) in a final volume of 50 μL was incubated on ice for 30 min in high salt buffer (25 mM 4-(2-hydroxyethyl)-1-piperazineethanesulfonic acid [HEPES], 2 M KCl, 1 mM EDTA [ethylenediamine tetraacetic acid], 10 mM DTT [1,4-dithiothreitol], 0.1 mg/mL BSA [bovine serum albumin], adjusted to pH 7.5). After core histones pre-incubation, 2 ng/μL of biotinylated DNA was added to the mixture at a final volume of 100 μL and further incubated on ice for 30 min. Then, the 100 μL mixture of DNA and core histones was put into a Slide-A-Lyzer device of 7000 MWCO (ThermoFisher) and dialyzed overnight against low salt buffer (25 mM HEPES, 50 mM KCl, 0.1 mM EDTA, pH 7.5). Reconstituted nucleosomes were diluted 20 times in low salt buffer before they were applied into the microfluidic chamber; otherwise, samples were stored at 4°C for up to three days. Under these assembly conditions, we found DNA molecules that would typically show from 0 to 6 unwrapping events during force-spectroscopy experiments.

Fluorescent labeling was performed by targeting histone amine groups of pre-assembled nucleosomes using Alexa Fluor 647 succinimidyl ester (ThermoFisher), following the previously reported strategy^20^. A mixture of ~2 ng/μL DNA with reconstituted nucleosomes, and 200 μM fluorophore reagent, was set in 25 mM HEPES, 50 mM KCl, 0.1 mM EDTA, 0.1 mg/mL BSA, 0.02% (v/v) Tween 20, 2% dimethyl sulfoxide (DMSO), pH 7.5. After one-hour incubation at room temperature, the reaction was diluted 20 times in low salt buffer and used in the microfluidic cell. Labeled nucleosomes were prepared fresh and discarded at the end of the day.

### Correlated optical tweezers and confocal fluorescence microscopy

A commercial set-up (LUMICKS) combining dual-trap optical tweezers, 3-color confocal fluorescence microscopy and microfluidics was used^53^. Two DNA constructs, bacteriophage λ DNA (48,502 bp; Roche), and linearized pKYB1 vector (8,370 bp; New England Biolabs), were used in this study. Both DNA constructs were end-biotinylated and used in combination with streptavidin-coated polystyrene beads (Spherotech) of 3.11 μm and 1.76 μm in diameter, with trap stiffness typically between 0.2 and 0.5 pN/nm. 2D fluorescence images were generated by scanning the confocal volume along the area of interest and collecting the emission by single-photon avalanche photodiodes (APDs). Pixel size was set to 100 nm and emission intensity units are given in kHz (counts•10^3^/s or photons•10^3^/s). Kymographs were constructed by collecting 1D scans along the DNA molecule over time at constant frequency, typically between 4 and 8 Hz. Experiments were performed in low salt buffer (25 mM HEPES, 50 mM KCl, 0.1 mM EDTA, pH 7.5).

### Formation of DNA-histone-chaperone complexes by OT and microfluidics

A motorised stage was used to control the position of a 5-channel microfluidic chamber, and thus rapidly change solution conditions. Individual λ-DNA molecules were isolated using OT and stretched up to 10 pN tension (~15.8 μm bead-to-bead distance) in low salt buffer solution. While keeping the trap-to-trap distance constant, to avoid the formation of DNA loops, DNA molecules were incubated with a solution of 100 nM of individual histones in low salt buffer for ~30 sec. In the case of H3, 10 mM DTT was added to avoid disulfide bond formation. Next, histone-DNA complexes were either brought back to the buffer solution, or to a solution of 5 nM chaperone in low salt buffer to study chaperone activities. OT-CFM experiments were performed for up to 5 min.

### Histone-DNA unbinding kinetics

Histone-DNA interaction was monitored by following the bead-to-bead distance, under different conditions. Bead-to-bead distances were determined by bright-field imaging of the beads in combination with a bead tracking algorithm. Distance was chosen over force readouts because our set-up relies on back focal plane interferometry^54^ to detect the forces being exerted on the beads. This makes force readouts very sensitive to changes in the light path, which can be severe for different regions of the microfluidic chamber.

The bead-to-bead distance was continuously monitored while introducing preformed histone-DNA complexes into either buffer, chaperone solutions, or a mixture of chaperone and C*c*. The increase in distance measured over time, towards bare DNA values, is then assumed to be related to the amount of histone bound (Fig. S5). Kinetic traces of 2 min were recorded for individual histone-DNA complexes under the different conditions reported. Averaged kinetic traces (Fig. 3, and S5) were generated from at least 5 different single-molecule traces, and fitted to an exponential function to extract the unbinding rates. Traces obtained in low salt buffer were all well described by a single exponential function. However, traces recorded in chaperone solutions showed deviations from a monotonic behavior; thus, a double exponential function was used for analysis. The errors associated with the reported off-rates represent SEM, obtained from the fits to individual single-molecule kinetic traces.

### Force-extension curves (FECs) proving chaperone shielding

FECs were performed on histone-DNA complexes after 5 min incubation in either low salt buffer or chaperone solutions, where trap-to-trap distance was not changed to keep DNA molecules extended. After incubation, curves were generated by rapidly approaching one of the traps at ~8 μm bead-to-bed distance (no tension applied), and then moving the trap at constant speed (500 nm/s) until reaching 50 pN of tension. Bare DNA curves were modeled by the extensible worm-like chain (eWLC) model^55^:

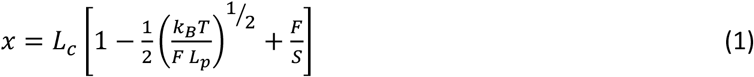

With *x*, extension; *F*, force; *L_c_*, contour length; *L_p_*, persistence length; *S*, elastic modulus; and *k_B_T*, Boltzmann constant times absolute temperature. The parameters that characterize the mechanical response of DNA were estimated experimentally from the eWLC fits to individual FEC of bare DNA molecules (N = 24) in low salt buffer, obtaining *L_c_* = 16.45 ± 0.01 μm, *L_p_* = 45 ± 1 nm, and *S* = 1690 ± 50 pN (mean ± SEM).

### Fluorescence imaging and fluorophore brightness estimation

Fluorescence images of fluorescently labeled chaperones were recorded at the rate of one image per min (image time ≈ 5 s) for up to 5 min, to monitor chaperone-histone interactions while minimizing photobleaching. Images at 0 min were always taken within the first 30 s of incubation, which includes the time needed to introduce the DNA-histone construct into the microfluidic channel where imaging is performed, containing either chaperone or chaperone-C*c* solutions. The 532 nm laser was set to ~1 μW power at objective, and pixel time of 0.5 ms. Kymographs were constructed by collecting 1D scans along the DNA at constant frequency, typically between 4 and 8 Hz. Mean fluorescence intensity (MFI) was calculated by extracting the mean number of photons per second. In the cases where individual binding events were identified, MFI was calculated for the individual traces considering a trace thickness of four pixels.

To estimate the count rate per fluorophore under our experimental conditions, a mixture of 10 nM H3, 10 nM H4, and 1 nM SET/TAF-Iβ in low salt buffer was applied to the microfluidic chamber. These conditions revealed individual binding events of SET/TAF-Iβ and DNA colocalization, which were well isolated and allowed fluorophore brightness determination. Excitation settings were kept constant, as described above. According to our labeling strategy and the dimeric nature of SET/TAF-Iβ, our analysis reported 11 ± 3 kHz per fluorophore and 21 ± 2 kHz per dimer (Fig. S3).

### Nucleosome unwrapping

Nucleosomes reconstituted on linearized pKYB1 molecules were isolated and brought to a solution of 1-2 nM chaperone without applying any tension to the tether. After an incubation of ~1 min, force-spectroscopy experiments were performed. FECs were generated by moving one of the traps at a constant speed of 20 nm/s between 1 and 32 pN. The changes in *L_c_* were obtained by fitting the eWLC model (equation 1) before and after any unwrapping event, while keeping *L_p_* and *S* constant and equal to the ones obtained for bare DNA. For pKYB1, we estimated *L_c_* = 2.818 ± 0.004 μm, *L_p_* = 50 ± 1 nm, and *S* = 1400 ± 40 pN (mean ± SEM; N = 20).

Time correlated fluorescence imaging was used to monitor fluorescently labeled histones and chaperones during the FECs. Kymographs were performed at 4 Hz (250 ms/line scan) and with an excitation laser of 532 nm and 638 nm, both set to ~1 μW power at objective. MFI of individual fluorescence traces were obtained as described above.

## Acknowledgments

We acknowledge funding through a Vidi grant of NWO (to WHR), and via a travel grant of the European ARBRE-Mobieu consortium (COST Action CA15126) (to ADQ). The authors are also grateful to the Spanish Ministry of Science and Innovation - Research Agency (PGC2018-096049-B-I00/FEDER), the Regional Government of Andalusia (US-1254317; US-1257019; P18-FR-3487; P18-HO-4091 -US/JUNTA/FEDER,UE and BIO198) and the Ramón Areces Foundation (2021-2024). We thank Gijs Wuite and Rifka Vlijm for critical feedback on the manuscript and discussions.

## Author Contributions

P.B. designed and performed the experiments, analyzed and interpreted the data, and wrote the initial draft of the paper. A.V.C., K.G.A., A.D.Q. and I.D.M. provided protein samples. All authors contributed to the interpretation of the results and to the final version of the manuscript. W.H.R. designed and supervised the study.

## Supplementary Figures

**Fig. S1.**
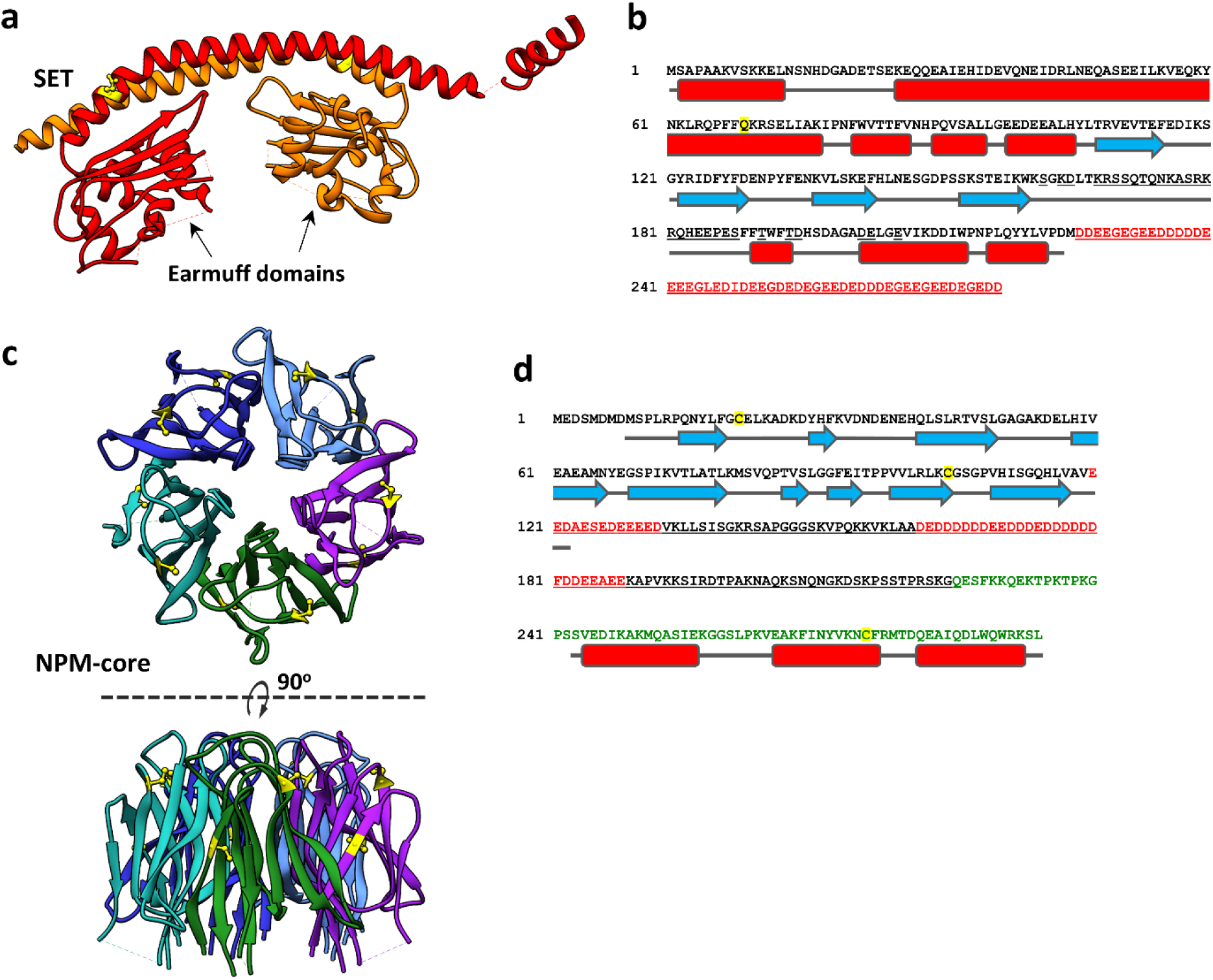
Structure of the histone chaperones SET/TAF-Iβ and NPM. **a**, Ribbon representation of SET/TAF-Iβ crystal structure with the individual monomers colored in red and orange, and the residue Q69 highlighted in yellow (disordered region is not shown). Each monomer contains an α-helix involved in the protein dimerization followed by an earmuff domain (PDB 2E50)^12^. **b**, Amino acid sequence of SET/TAF-Iβ showing its secondary structure elements: α-helices (red) and β-strands (cyan). The disordered acidic region is colored in red and the residue Q69, mutated to Cys for fluorescent labeling, is highlighted in yellow. Residues involved in histone interactions are underlined. **c**, Ribbon representation of the pentameric NPM-core, amino acids sequence 1-122 (PDB 2P1B)^13^, with Cys residues highlighted in yellow. View of the five-fold symmetry axis (upper panel), and side view (lower panel). **d**, Sequence of NPM showing its secondary structure elements: α-helices (red) and β-strands (cyan), according to^13, 56^, with Cys residues highlighted in yellow. The region of the C-terminal domain identified for the interaction of NPM with DNA is colored in green^28^. The disordered acidic region is colored in red and residues involved in histone interactions are underlined. Molecular models were generated with UCSF Chimera^57^.

**Fig. S2.**
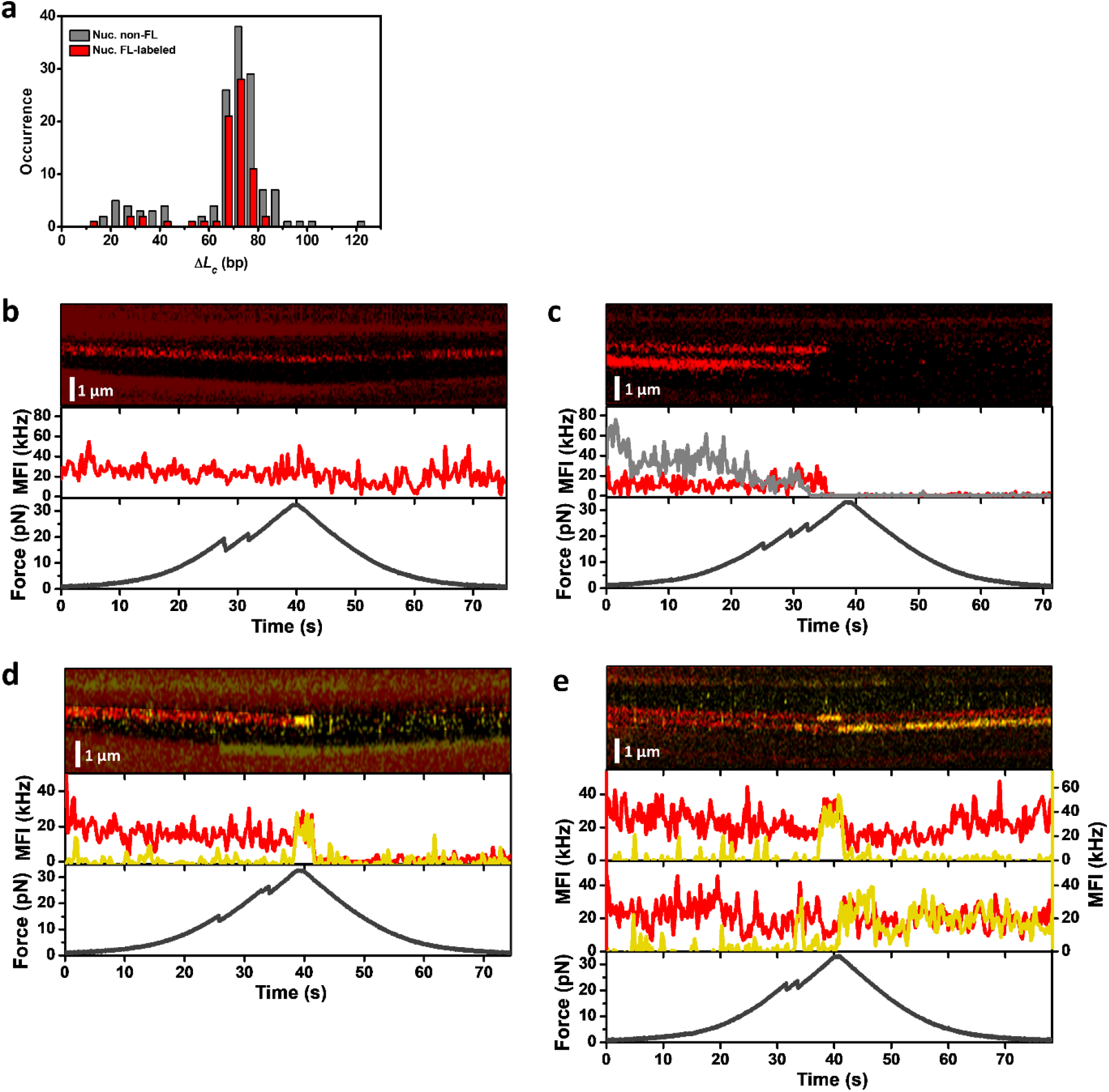
Correlated FEC and fluorescence imaging of fluorescently labeled histones and chaperones. **a,** Distributions of the *L_c_* changes obtained from nucleosome unwrapping experiments with non-labeled histones (grey; N = 138) and with fluorescently labeled histones (red; N = 71). **b**, Example of fluorescently labeled histones that remain bound after nucleosome unwrapping was detected. **c**, Example showing histone signal disappearing soon after unwrapping events were found. Middle panel shows MFI of the most upper trace (red) and lower trace (grey) of the kymograph. **d**, Example of histone (red) eviction by the chaperone SET/TAF-Iβ (yellow) in which the fluorescence signal of both histone and chaperone fall to background levels upon chaperone unbinding. **e**, Correlated FEC and fluorescence imaging showing two examples in which the histone signal (red) does not change upon unwrapping or chaperone binding (NPM; yellow).

**Fig. S3.**
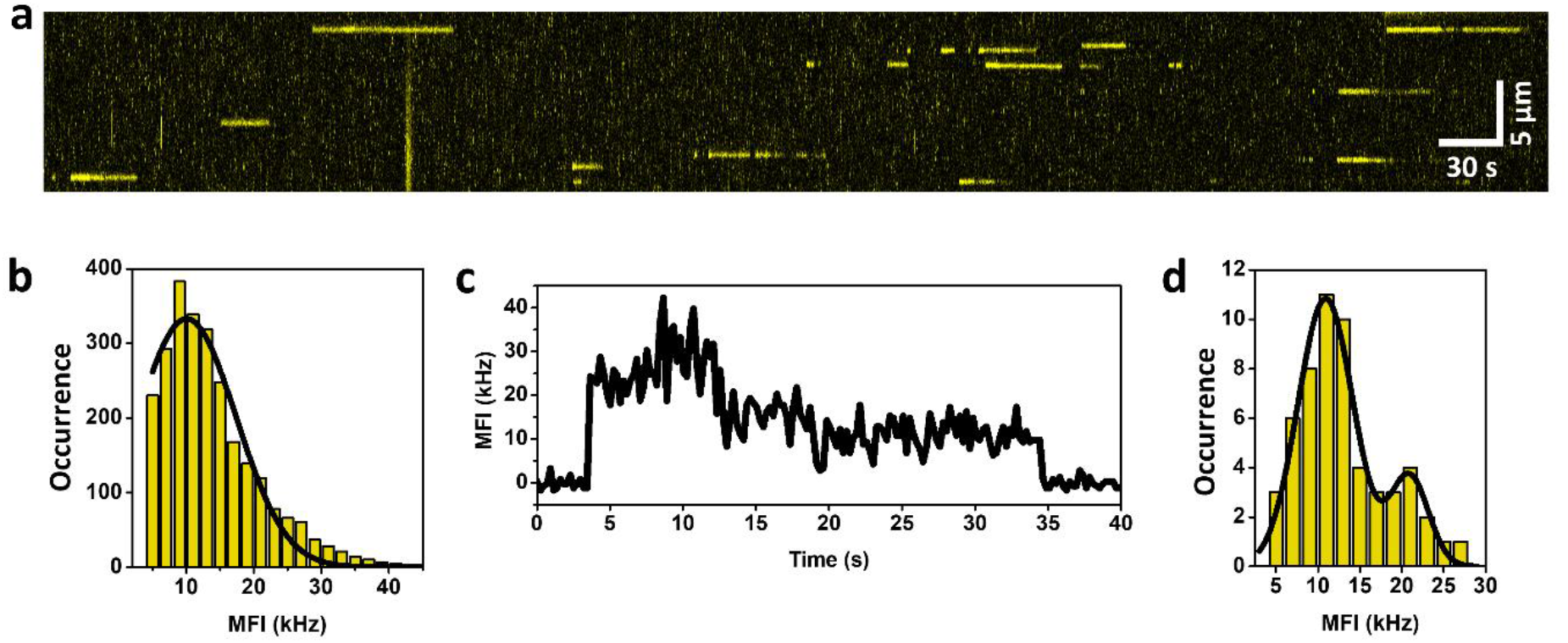
Fluorophore brightness. **a**, Kymograph recorded at 4.4 Hz (228ms/line) in the presence of 10 nM H3, 10 nM H4, and 1 nM of fluorescently labeled SET/TAF-Iβ (Alexa Fluor 532) in low salt buffer. **b**, Yellow bars show the MFI distribution obtained from 33 individual traces after background correction. MFI is calculated for every time point, averaging the intensity of four pixels in the distance axis, of the individual fluorescence traces. The black solid line shows the fit to a single Gaussian function. Gaussian fit reported 10 ± 7 kHz (center ± SD). **c**, Representative MFI *versus* time trace showing two discrete levels of intensity due to photobleaching; consistent with the labeling of dimeric SET/TAF-Iβ that produced the attachment of two fluorophores per dimer on average (see methods). **d**, MFI distribution constructed from the averaging of individual traces into two discrete intensity levels, high and low. Occurrence represents the total number of time-regions with constant MFI. Fit to a double Gaussian reported 11 ± 3 kHz per fluorophore and 21 ± 2 kHz per dimer (center ± SD).

**Fig. S4.**
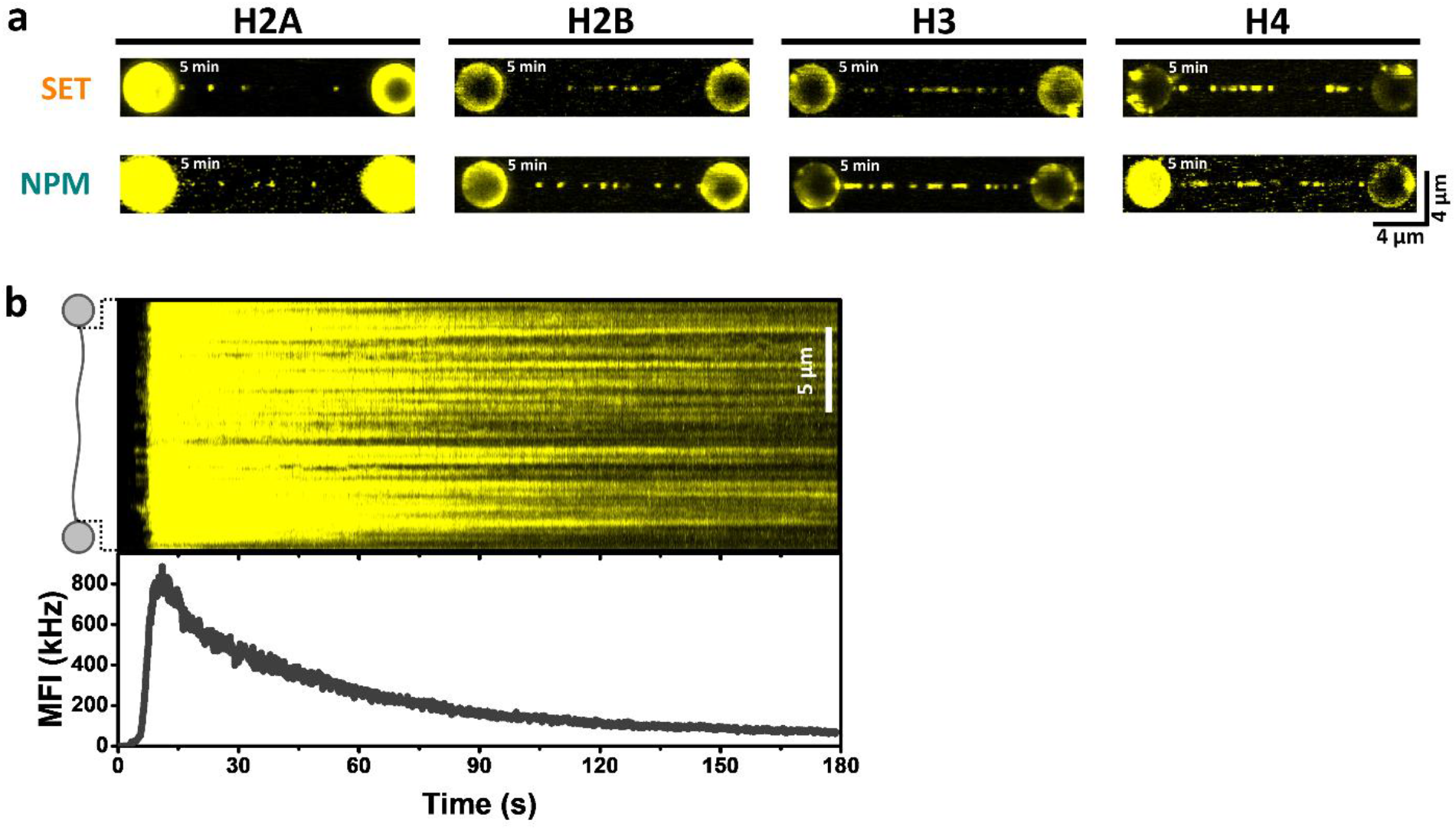
Visualization of histone recognition by fluorescence imaging of histone chaperones. **a**, Fluorescence images of 5 nM SET/TAF-Iβ (top row) and 5 nM NPM (bottom row) on DNA, after incubating the DNA-histone complex for 5 min with one type of histones (either H2A, H2B, H3, or H4, from left to right). The DNA molecules were stretched to a fixed distance, corresponding to 10 pN tension, and were then incubated in a solution of individual histones at 100 nM concentration for ~30 s. Next, histone-DNA complexes were brought to a solution of fluorescently labeled chaperone and observed for up to 5 min at the rate of one image per minute (image time ≈ 5 s). **b**, Upper panel, kymograph of a H2B-DNA complex entering the NPM solution recorded at 7.6 Hz (131 ms/line) through continuous imaging. Lower panel, mean fluorescence intensity (MFI) per line scan over time.

**Fig. S5.**
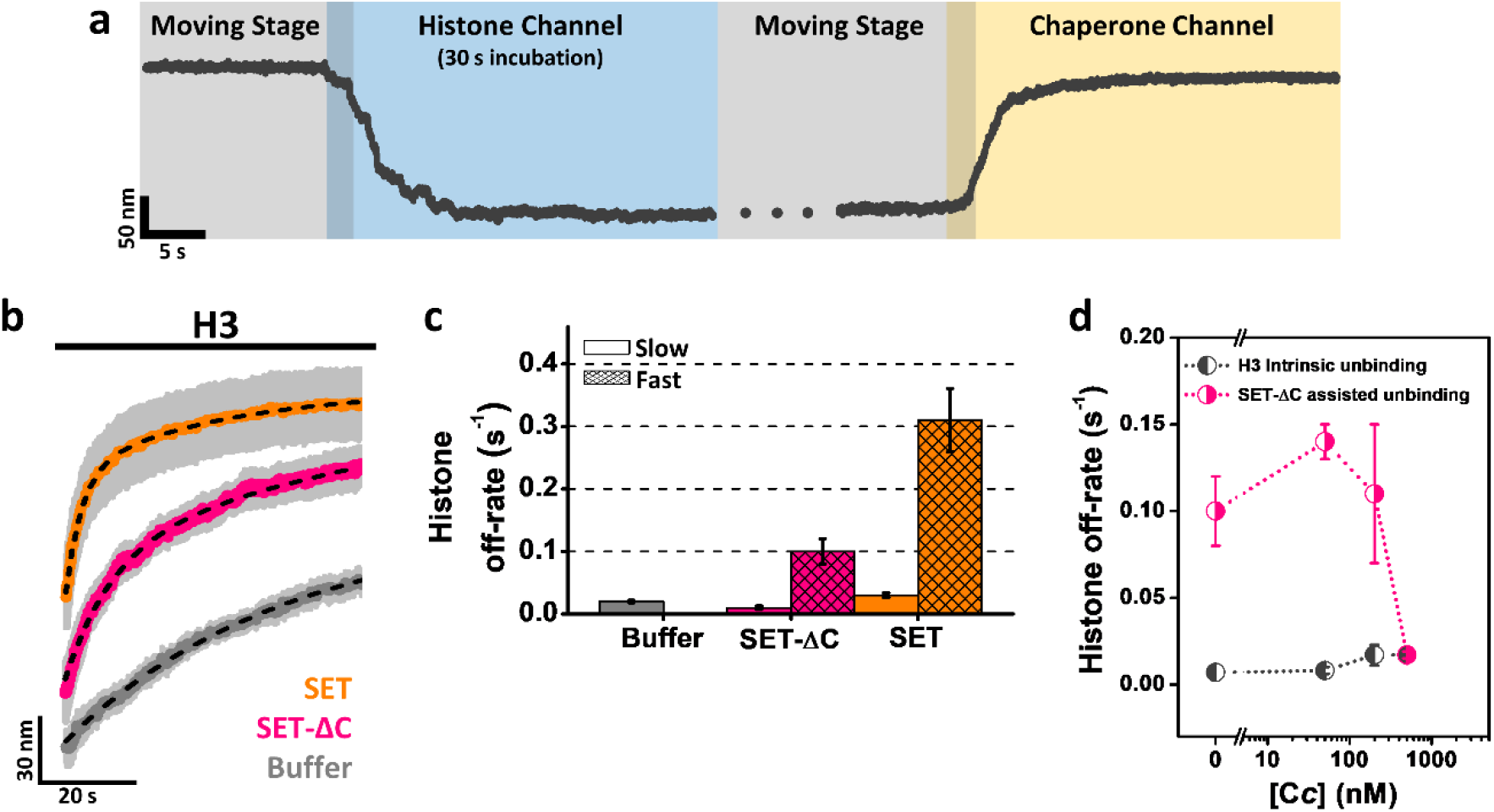
Binding kinetics of histone-DNA interactions mediated by histone chaperones. **a**, Example of a full kinetic experiment to measure histone eviction by following the changes in bead-to-bead distance. DNA molecules were stretched to a bead-to-bead distance of ~15.8 μm (~10 pN) in the buffer solution. Then, the DNA molecule is brought into a solution of individual histones at 100 nM concentration by moving the stage of the microfluidic cell holder. Incubation took place for ~30 s, where a rapid decrease in distance, DNA condensation, was observed (light blue). Next, histone-DNA complexes were brought to a solution of 5 nM chaperone (light yellow). Off-rates were calculated from 2 min traces of the bead-to-bead distance during chaperone incubation. **b**, Averaged kinetic traces of H3-DNA complexes during their incubation against buffer (grey), 5 nM SET/TAF-Iβ (orange), and 5 nM SET/TAF-Iβ-ΔC (pink). Averaged curves were generated from individual traces (N ≥ 5) and fitted to either a single or double exponential function (black dashed lines). Grey shades represent SEM. **c**, Off-rates obtained from the fits presented in (b). Error bars represent SEM, calculated from the fits to the individual kinetic traces. **d**, Off-rates at different C*c* concentrations (0, 50, 200, and 500 nM) obtained from H3-DNA complexes that were brought to a mixture solution of 5 nM SET/TAF-Iβ-ΔC and C*c*. The slow rate is depicted in dark grey, while the fast rate is colored in pink. Error bars represent SEM.

**Fig. S6.**
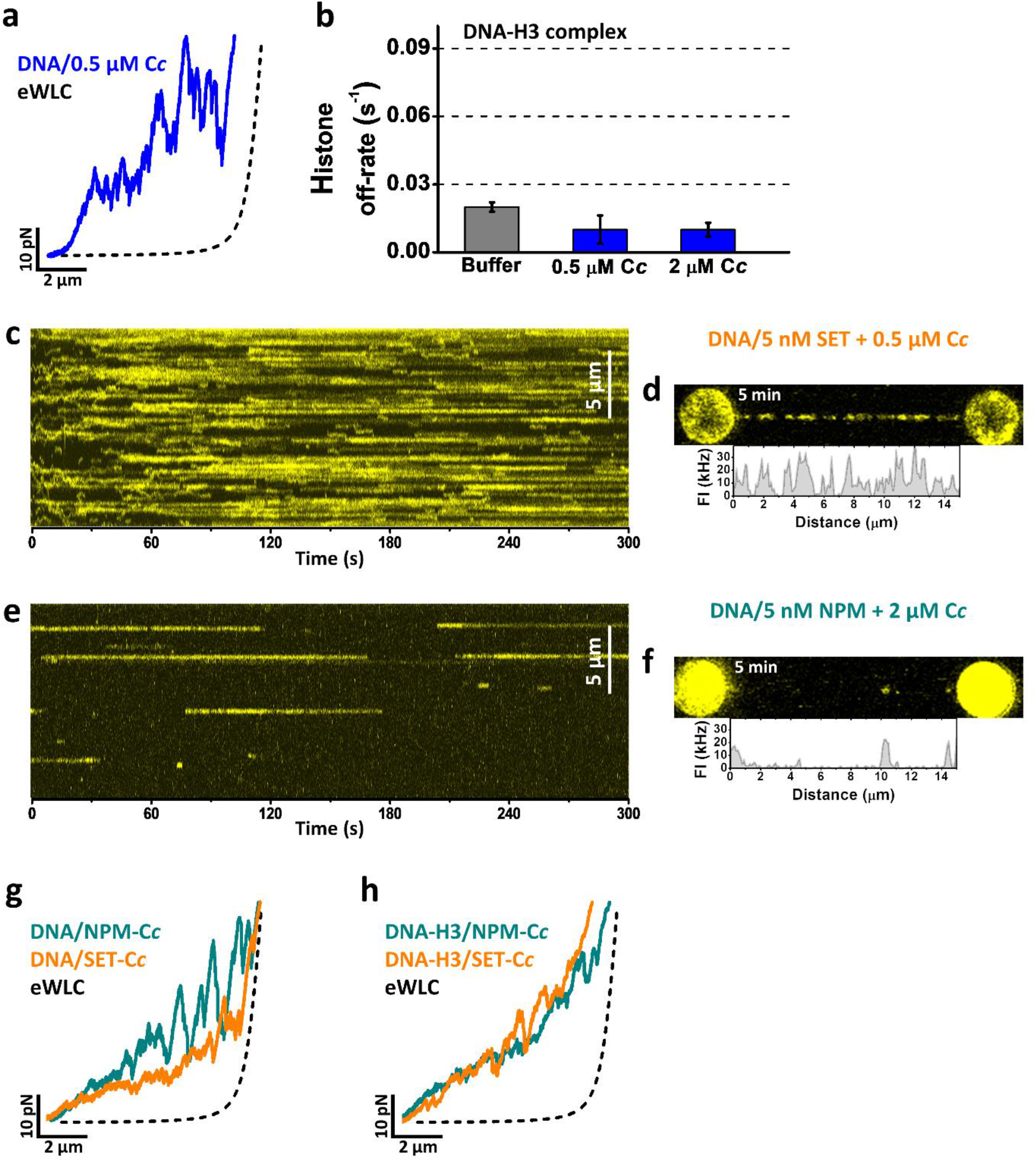
C*c*-DNA interactions in the presence and absence of chaperones. **a**, Representative FEC of DNA in a solution of 0.5 μM C*c* (blue). **b**, Off-rates obtained from H3-DNA complexes in a solution of either 0.5 μM or 2 μM C*c* (blue). The intrinsic unbinding rate measure for H3 is shown in grey for comparison. Error bars represent SEM, calculated from the fits of individual kinetic traces. **c**, Kymograph of a DNA molecule in a solution of 5 nM SET/TAF-Iβ and 0.5 μM C*c*. **d**, Fluorescence image after 5 min incubation under the same conditions described in (c). **e**, Kymograph of a DNA molecule in a solution of 5 nM NPM and 2 μM C*c*. **f**, Fluorescence image after 5 min incubation under the same conditions described in (e). **g**, Representative FECs of DNA in a mixture solution of 5 nM SET/TAF-Iβ and 0.5 μM C*c* (orange), and 5 nM NPM and 2 μM C*c* (dark green). **h**, FECs of H3-DNA complexes in the presence of 5 nM SET/TAF-Iβ and 0.5 μM C*c* (orange), and 5 nM NPM and 2 μM C*c* (dark green).

